# The chromatin-binding domain of Ki-67 together with p53 protects human chromosomes from mitotic damage

**DOI:** 10.1101/2020.10.16.342352

**Authors:** Osama Garwain, Xiaoming Sun, Divya Ramalingam Iyer, Rui Li, Lihua Julie Zhu, Paul D. Kaufman

## Abstract

Vertebrate mammals express a protein called Ki-67 which is most widely known as a clinically useful marker of highly proliferative cells. Previous studies of human cells indicated that acute depletion of Ki-67 can elicit a delay at the G1/S boundary of the cell cycle, dependent on induction of the checkpoint protein p21. Consistent with those observations, we show here that acute Ki-67 depletion causes hallmarks of DNA damage, and the damage occurs even in the absence of checkpoint signaling. This damage is not observed in cells traversing S phase but is instead robustly detected in mitotic cells. The C-terminal chromatin binding domain of Ki-67 is necessary and sufficient to protect cells from this damage. We also observe synergistic effects when Ki-67 and p53 are simultaneously depleted, resulting in increased levels of chromosome bridges at anaphase, followed by the appearance of micronuclei. Therefore, these studies identify the C-terminus of Ki-67 as an important module for genome stability.

## Introduction

Mammalian proliferation antigen Ki-67 has well-established clinical significance because of its utility as a marker for aggressive tumor cells (reviewed in (1)). Ki-67 is rapidly degraded during the G1 phase of the cell cycle, so quiescent or slowly growing cells that have long G1 phases generally have low levels of Ki-67 (2). In contrast, rapidly growing cells, including tumor cells that lack checkpoint controls, often have short G1 phases and thereby display high steady-state Ki-67 levels (3). High Ki-67 protein levels are correlated with the severity of many types of tumors, and are a strongly predictive of poor outcomes in meta-analyses of clinical cancer data (4–7). As we describe below, Ki-67 is critical for maintaining several aspects of chromosome structural integrity. Therefore, molecular exploration of these functions is crucial to understanding how Ki-67 contributes to tumor biology.

Clues regarding the multiple functions of Ki-67 come from its dramatic relocalization across the cell cycle (1, 8, 9). In interphase cells, Ki-67 is localized to the nucleolus and is required for efficient localization of heterochromatin to the nucleolar periphery (10–12). Depletion of Ki-67 also alters the focal accumulation of the heterochromatic histone modification H3K9me3 (11) and the modification and localization of nucleolus-associated inactive X chromosomes (13). After interphase, Ki-67 localization dramatically changes when it becomes heavily phosphorylated by CDK1 during mitotic entry (14, 15). Ki-67 coats mitotic chromosomes, serving as a fundamental component of the perichromosomal layer (PCL; (10, 16); reviewed in (1, 17, 18)). The PCL is the mitotic repository of the abundant ribonucleoprotein complexes that inhabit the nucleolus during interphase (18, 19). Without Ki-67, the PCL is absent, dispersing these components (10, 16), causing imbalanced inheritance of nucleolar material in daughter cells (10). During mitosis, Ki-67 has additional key functions: it is required for maintenance of spatially separated chromosome arms (20, 21) and also for the clustering of chromosomes prior to nuclear envelope reformation, thereby preventing retention of cytoplasmic material when nuclei reassemble after mitosis (22). Tethered to chromatin, the long and highly charged Ki-67 protein serves as an electrostatic repellant that prevents chromosome arms from clumping (20, 21). This pathway is distinct from the contribution of condensin proteins to mitotic chromosome structure, because co-depletion of both Ki-67 and condensin results in synergistic loss of nearly all mitotic chromosome structure (23). In sum, Ki-67 shapes chromosome architecture in both interphase and mitotic cells.

In our previous studies, we had observed that acute depletion of Ki-67 causes a delay in cell cycle progression at the G1/S transition of the cell cycle in checkpoint-proficient cells (13). This delay is accompanied by induction of the cyclin-dependent kinase inhibitor p21, which is required for the delay (13). p21 is a transcriptional target of the tumor suppressor protein p53 (24, 25), a critical activator of the transcriptional response to DNA damage. These observations led us to test whether Ki-67 protects cells from DNA damage, and whether p53 has a role. We show here that acute depletion of Ki-67 results in DNA damage as evidenced by increased levels of modified histone γH2AX and focal accumulation of repair signalling protein 53BP1. This damage accumulates during mitotic progression in both in cells that display a checkpoint response to Ki-67 depletion and in cells that do not, indicating that the role of Ki-67 in genome stability is independent of a transcriptional response to damage. We demonstrate that the C-terminal chromatin binding domain of Ki-67 is necessary and sufficient to protect cells from this damage. These data define a novel type of genome protection molecule, and indicate that this activity is distinct from Ki-67’s role in maintaining distinct chromosome arm structures, which requires additional parts of the protein to provide sufficient electrostatic repulsion (20). We also show that loss of Ki-67 causes greater defects in the absence of tumor suppressor protein p53, including a large increase in the numbers of anaphase bridges that appear during the first mitosis after acute Ki-67 depletion. Subsequently, cells lacking Ki-67 and p53 frequently display micronuclei that lose lamin A protein from their periphery over time, a hallmark of membrane disruption associated with genome rearrangements (26, 27). In sum, these data indicate that Ki-67 physically protects chromosomes from damage during mitosis.

## Results

### Ki-67 protects cells from DNA damage regardless of G1/S checkpoint status

To test whether acute depletion of Ki-67 causes DNA damage, we first analyzed cells by immunofluorescence using antibodies that recognize γH2AX, a phosphorylated histone isoform that is a classical marker of DNA strand breaks (28). We initially analyzed hTERT-RPE1 cells, a diploid, checkpoint-profiecient human cell line, which display a p21-dependent transcriptional program in response to Ki-67 depletion (13). Using the same si-RNA duplex we had validated previously in RPE-1 cells (13), we tested acute depletion of Ki-67 in an asynchronous cell population, thereby sampling all cell cycle positions. We observed that depletion of Ki-67 significantly increased the γH2AX signals (Fig. 1A), and that the magnitude of this effect was amplified in the presence of the DNA strand-breaking reagent phleomycin (Fig. 1B; (29)). These data suggested that DNA damage occurs upon acute depletion of Ki-67.

**Figure 1.**
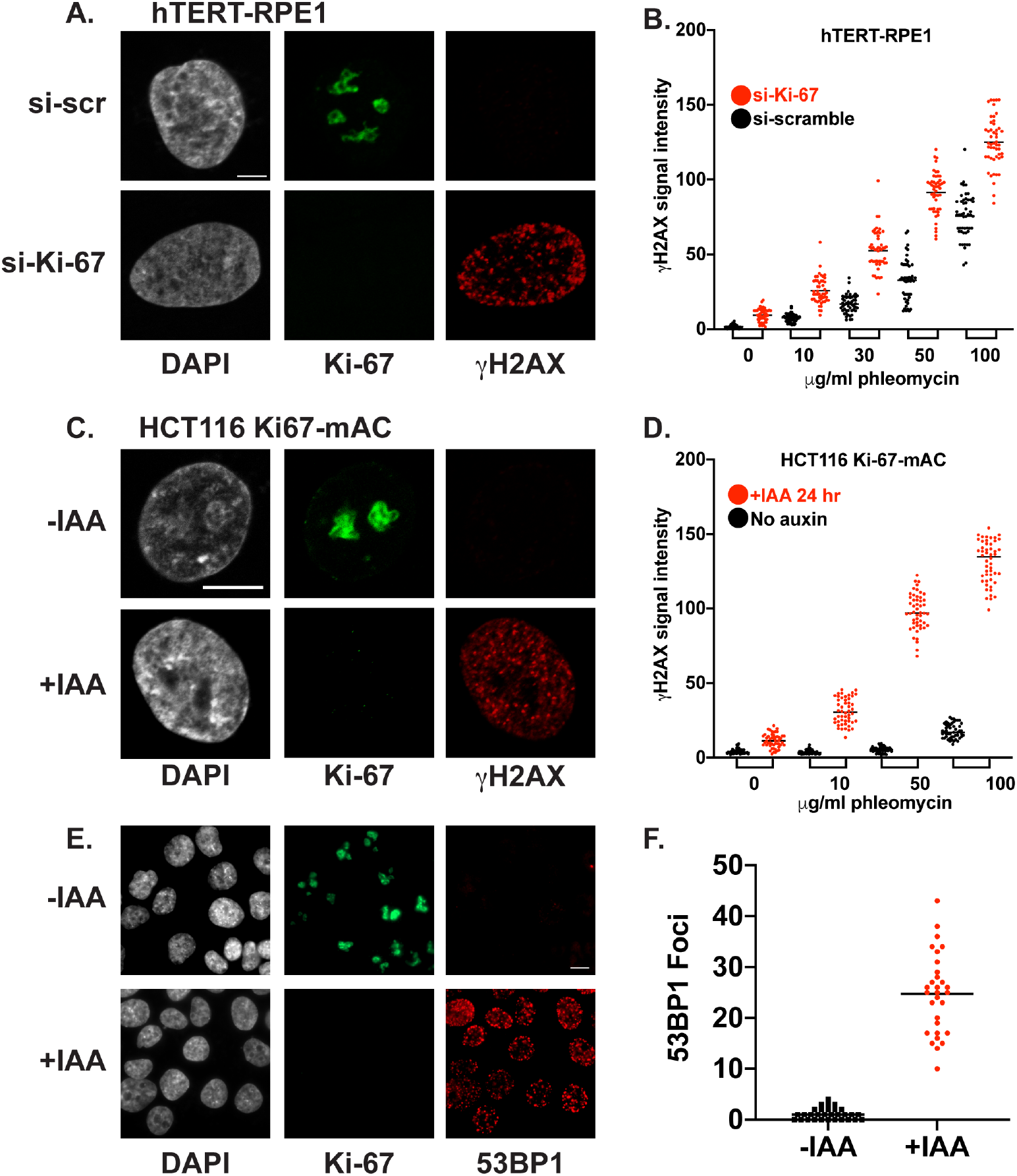
γH2AX and 53BP1 staining in Ki-67-depleted cells. Scale bars are 5 μm and magnifications are the same within each panel. (A) hTERT-RPE1 cells were treated with si-scr or si-Ki-67 as indicated for 72 hours. DNA was stained with DAPI (grey), and Ki-67 (green) and γH2AX (red) were detected by immunofluorescence (IF). (B) hTERT-RPE1 cells were treated as in (A), in the presence or absence of the indicated amounts of phleomycin. γH2AX signals from 50 cells of each population are graphed, with mean values indicated by a black line. Ki-67 depletion significantly increased γH2AX signals for all doses of phleomycin (p < 0.0001, Welch’s t-test). (C) HCT116 Ki-67-mAC cells (hereafter called HCT116 cells) were either untreated (-IAA) or treated with indole acetic acid (+IAA) for 24 hours to deplete Ki-67, and analyzed as in panel (A), except that Ki-67 was detected via fluorescence of the mClover tag. (D) Quantitation of γH2AX in HCT116 cells, as in panel (B). Ki-67 depletion significantly increased γH2AX signals at all doses of phleomycin (p < 0.0001, Welch’s t-test). (E) HCT116 cells were treated as indicated and prepared for IF with antibodies recognizing 53BP1 (red). Green channel fluorescence detected mClover-tagged Ki-67. (F) 53BP1 foci were counted in thirty cells in each population from the experiment in panel (E).

Our previous studies showed that upon acute depletion of Ki-67, RPE-1 cells display a transient cell cycle delay at the G1/S boundary, accompanied by p21-dependent down-regulation of many S phase-related transcripts (13). Therefore, we considered the possibility that the transcriptional response to Ki-67 depletion was indirectly causing DNA damage. To test this idea, we analyzed a cell line that lacks this transcriptional response, colon cancer HCT116 cells. Specifically, we used a derivative of this well-studied line in which homozygous in-frame insertions encode Ki-67 tagged with a fluorescent mClover protein and an auxin-inducible degron (21), which allows rapid depletion of detectable Ki-67 upon addition of auxin (indole acetic acid, IAA). We confirmed that these HCT116-Ki-67-mAC cells, (henceforth HCT116 for brevity), unlike RPE-1 cells but similar to virally-transformed HeLa cells, do not induce p21 mRNA levels in response to Ki-67 depletion (Fig. S1A). Furthermore, RNA-seq analysis of HCT116 cells detected an extremely limited genome-wide transcriptional response to acute Ki-67 depletion (Supplemental Figure S1B-D; Supplemental Table 1). Nevertheless, after 24 hours of auxin-mediated depletion of Ki-67, γH2AX signals were significantly increased in these cells (Fig. 1C-D). Together, these data indicate that Ki-67 has a role in genome stability in multiple cell types.

These data also suggested that the contribution of Ki-67 to genomic stability is independent of the p21-mediated transcriptional response. To test this directly, we codepleted p21 and Ki-67. We observed that depletion of p21 did not significantly change the levels of γH2AX generated either in the absense or presence of Ki-67, even when the assay was sensitized by the presence of phleomycin (Supplemental Figure S2A-C). These data are consistent with our finding that the damage caused by depletion of Ki-67 does not depend on checkpoint signaling (Figure 1). We therefore hypothesized that Ki-67 has a direct role in protecting the genome from damage.

### 53BP1 foci are formed upon depletion of Ki-67

Our discovery of DNA damage upon Ki-67 depletion was surprising because a previous study discounted the possibility of a contribution to DNA protection by Ki-67, because no 53BP1 foci were detected in cells lacking Ki-67 (10). 53BP1 forms large foci at sites of DNA damage and is instrumental in regulating the choices between homologous recombination and endjoining repair pathways (30–33). Because of our observation of robust γH2AX signals, we re-examined whether Ki-67 depletion results in 53BP1 focus formation. Indeed, we find that it does (Fig. 1D-E), consistent with Ki-67 having an important role in genome protection. These 53BP1 foci were frequently overlapping γH2AX foci, consistent with the documented direct recruitment of 53BP1 by γH2AX (34). 53BP1 was not detected in mitotic cells regardless of the amount of DNA damage (Supplemental Figure S2D), consistent with previous experiments demonstrating that 53BP1 foci are not present in mitotic cells (35, 36).

### Ki-67 protects chromosomes during mitosis

In Figure 1, asynchronous populations were analyzed, so that cells across all points of the cell cycle had experienced the absence of Ki-67. We next sought to determine whether the role of Ki-67 in protection of chromosomes from damage might be important at specific times during the cell cycle. We first tested whether damage occurred during S phase. To do this, we pulse-labeled asynchronous HCT116 cell cultures with the modified deoxynucleotide EdU for twenty minutes prior to fixation and click-chemistry staining to detect cells that synthesized DNA during the pulse. The control population that was untreated with auxin did not display strong γH2AX signals, either in the EdU-positive subpopulation that had been in S phase, nor in the mitotic subpopulation that displayed chromosome condensation (Fig. 2A). In the IAA-treated population, we observed that the S phase subpopulation also lacked strong γH2AX signals. In contrast, the mitotic auxin-treated cells displayed robust γH2AX staining (Fig. 2B). These data suggested that mitosis is a particularly important time for genome protection by Ki-67.

**Figure 2.**
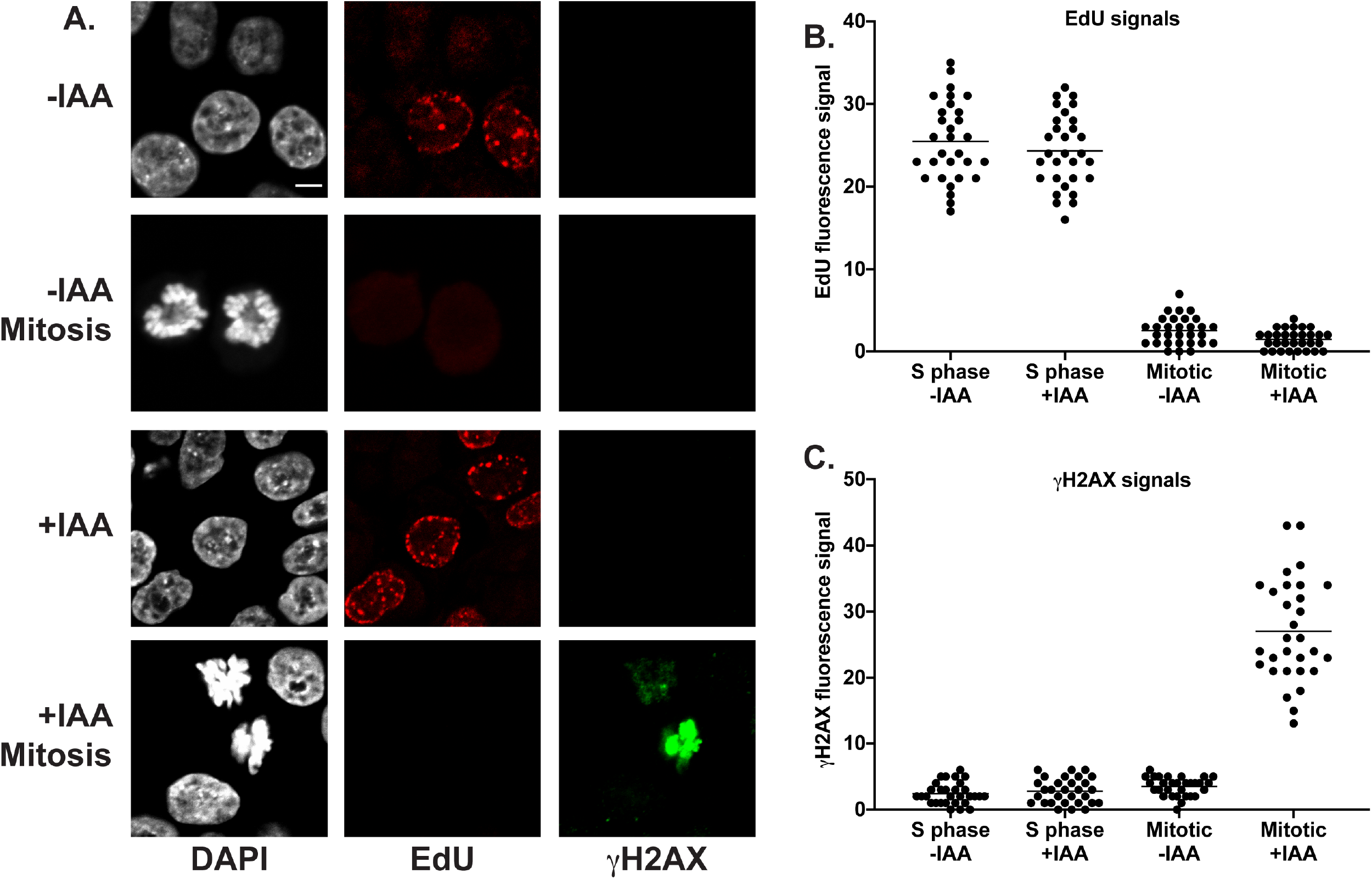
Ki-67 depletion causes damage in mitotic cells. (A) HCT116 cells were treated or untreated with IAA for five hours as indicated, and pulse-labeled with EdU (red) thirty minutes prior to fixation for IF detection of γH2AX (green). Mitotic cells in these asynchronous populations are indicated separately. Scale bar is 5 μm. (B) EdU signals in thirty cells of each of the indicated populations were quantified. “S phase” cells displayed visible EdU signals, “mitotic” cells were those that displayed condensed chromosomes. (C) The γH2AX signals in the same cells analyzed in panel B are displayed.

To test this more directly, we synchronized HCT116 cells at the G2/M phase border with the CDK1 inhibitor RO-3306 (37), depleted Ki-67 using auxin, and then measured damage upon release. In cells depleted of Ki-67, but not in control untreated cells, damage increased within minutes, concurrent with progression through mitosis (Figure 3). These data suggested that Ki-67 has a particularly important role in genome protection during mitosis. Furthermore, the rapid appearance of damage in these synchronized cell experiments suggest that this damage is unlikely to result from indirect transcriptional effects, consistent with the limited transcriptional effects of Ki-67 depletion in HCT116 cells (Supplemental Fig. S1). Together, these data supported the idea that Ki-67 has a direct role in protecting chromosomes.

**Figure 3.**
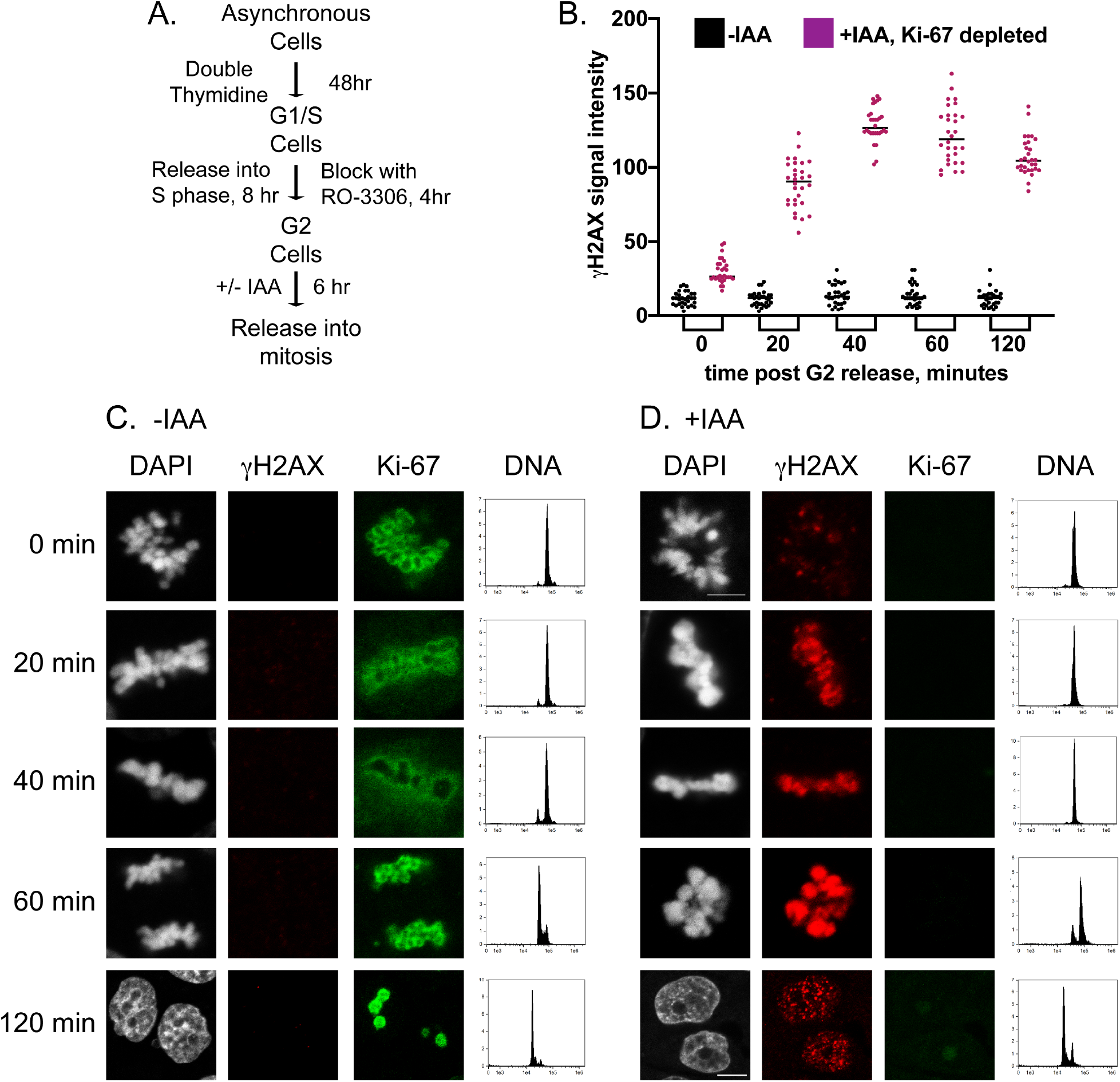
Damage occurs during mitotic progression in Ki-67-depleted cells. (A) Scheme for cell synchronization. (B) γH2AX signal intensity was measured in thirty cells in each of the control (-IAA) and auxin-treated (+IAA) populations from the indicated time points after release from RO-3306 arrest. (C-D) IF analysis of individual cells from the indicated time points in the (C) −IAA and (D) +IAA experiments. DAPI (grey), γH2AX (red) and Ki-67-mClover (green) are shown. Panels on the right show FACS analysis of the DNA content of DAPI-stained cells treated in the same manner as cells analyzed by IF. Scale bars are 5 μm, with the same magnification used for the 0-60-minute time points. The fields displaying cells from the 120-minute time point are slightly larger to capture two cells.

### The C-terminal chromatin-binding domain of Ki-67 is necessary and sufficient for protecting chromosomes

To determine whether specific domains of Ki-67 protect chromosomes from damage, we used HCT116-Ki-67-mAC cells to estabish an assay for Ki-67 transgenes. We transiently transfected plasmids encoding various GFP-tagged Ki-67 fragments (15) into asynchronous cell populations. Twelve hours later, we added auxin (IAA) and continued incubation for 24 hours to degrade endogenous Ki-67 and allow accumulation of damage, and then measured γH2AX and GFP signals by FACS and immunofluorescence analyses. We first confirmed that in untransfected cells, low levels of γH2AX staining (right hand side of the graph, percentage indicated above) were detected in more than 90% of the cells in the absence of auxin treatment (-IAA; Figure 4A). Conversely, high levels of damage were observed upon auxin treatment, with fewer than 10% of cells displaying γH2AX levels similar to the untreated population (+IAA, Figure 4A). We also noted that GFP signals were greatly decreased by auxin treatment, indicating efficient degron-mediated destruction of the endogenous mClover-tagged Ki-67 protein (Figure 4A). We then analyzed transfected cells treated with IAA. In empty vector (EGFP)-transfected cells, most cells displayed high GFP levels, but few undamaged cells were observed (Figure 4B). In contrast, transfection of a wild-type Ki-67 transgene also increased GFP levels but prevented most cells from displaying high levels of damage (Fig. 4B). Therefore, this assay was able to detect chromosome protection by Ki-67 transgenes. We then tested a series of deletion constructs (Fig. 4C), and observed that all constructs encoding the C-terminal chromatin-binding domain protected cells from elevated γH2AX levels, and all constructs that lack this domain did not (Fig. 4D-E). We note that in these experiments, transgenes provided the expected protein localization (15): during the critical mitotic period, Ki-67 derivatives containing the C-terminal chromatin-binding domain robustly coat the chromosomes, and those that lack this domain do not (Supplemental Figure S3). We conclude that the C-terminus of Ki-67, which is the chromosome binding domain (15, 20, 38), is necessary and sufficient for genome protection.

**Figure 4.**
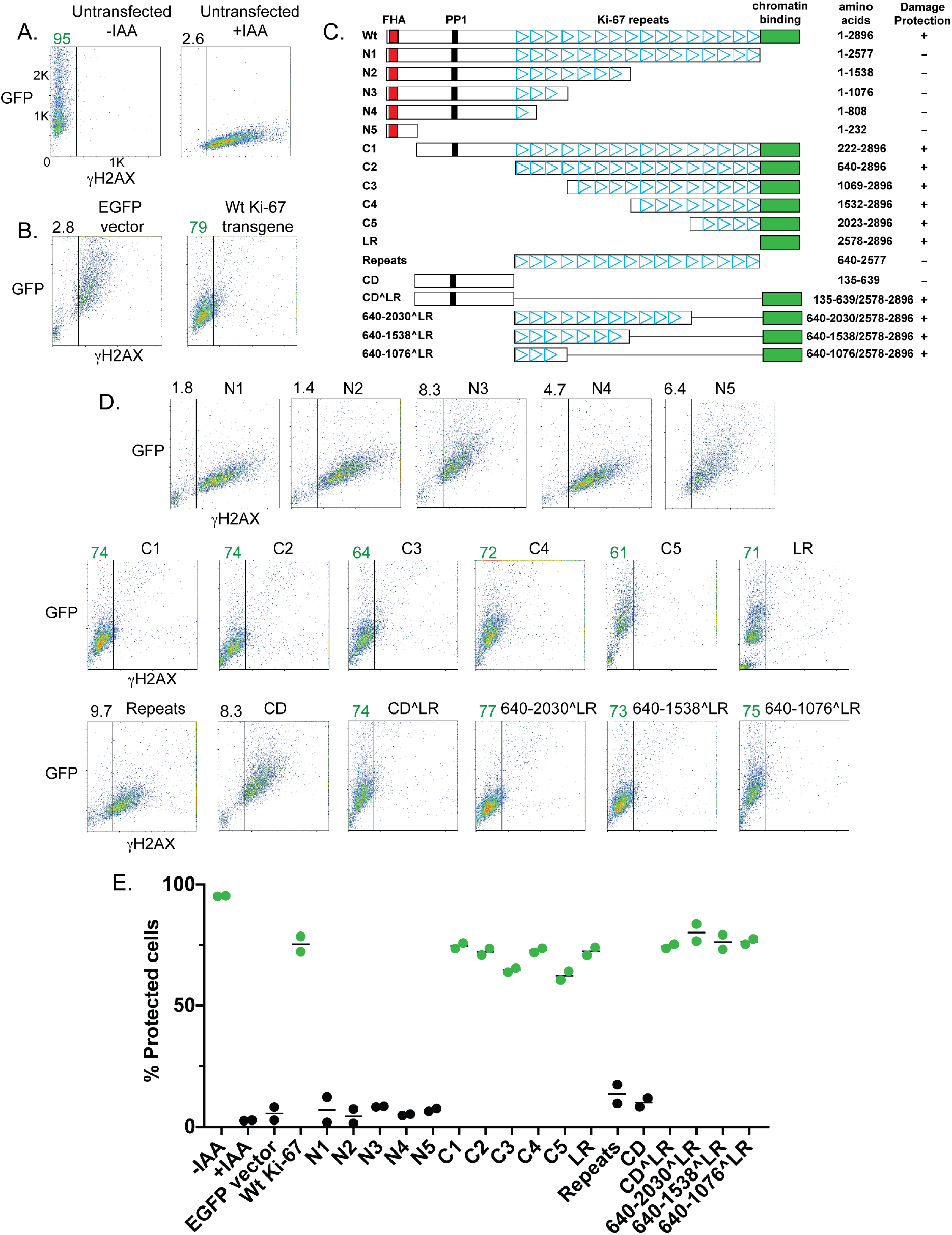
The C-terminal domain of Ki-67 protects cells from damage. (A) 2D FACS analysis of the HCT116-Ki-67-mAC cells, with GFP channel intensity on the y-axis and γH2AX signal intensity on the x-axis. Untransfected cells were either untreated (-IAA) or treated with auxin (+IAA). IAA treatment resulted in degradation of Ki-67-mClover, causing the loss of cells with low γH2AX signal intensity as well as fluorescence in the GFP channel. The percentages of cells with low γH2AX signal intensity (to the left of the vertical black line) are shown at the upper left corner. (B) Control experiments. Cells were transfected with the indicated constructs, treated with IAA, and analyzed by 2D FACS as above. Transfection of the empty vector (pEGFP-C1) restored GFP fluorescence to the majority of cells, but most cells displayed elevated γH2AX signal intensity. In contrast, transfection of a plasmid encoding GFP fused to a full-length Ki-67 cDNA resulted in most cells having reduced γH2AX signal intensity. (C) Schematic of transfected constructs. The FHA domain, the protein phosphatase 1-binding domain (PP1), the internal repeats (triangles) and the C-terminal chromatin-binding “LR” domain (green) are indicated. (D) 2D FACS analysis of cells transfected with the indicated constructs. Constructs that resulted in the majority of cells displaying low γH2AX signal intensity (“protected”) are tabulated in panel (C). (E) Summary of data from biological replicate experiments. Treatments yielding a majority of “protected” cells are colored in green.

### Synergies upon co-depletion of p53

In contrast to the effects of p21 depletion (13) (Figure S2), we observed a dramatic loss of viability in hTERT-RPE1 cells when Ki-67 and p53 were depleted simultaneously (Fig. 5A-B). We also observed synergy with p53 depletion in HCT116 cells, although in ths case reduced proliferation rather than lethality was observed (Fig. 5C). Furthermore, co-depletion of Ki-67 and p53 caused robust damage during synchronous progression through mitosis (Figure 5D-F), resulting in greater γH2AX phosphorylation signals than observed upon depletion of either protein alone (Figures 5D and 1D). To determine whether the same domain of Ki-67 protects cells in the absence of p53, we repeated the previous transgene complementation assay in HCT116 cells treated with si-p53 (Supplemental Figure S4). Again, the C-terminal LR domain was necessary and sufficient for protection, wtih the magnitude of the γH2AX signals greater in the absence of p53 (Supplemental Figure S4). We confirmed the synergistic damage phenotypes using a second siRNA that targets p53 (Supplemental Figure S5A-D). Together, these data indicated that the genomic damage caused by the lack of the Ki-67 C-terminal domain is magnified in the absence of p53.

**Figure 5.**
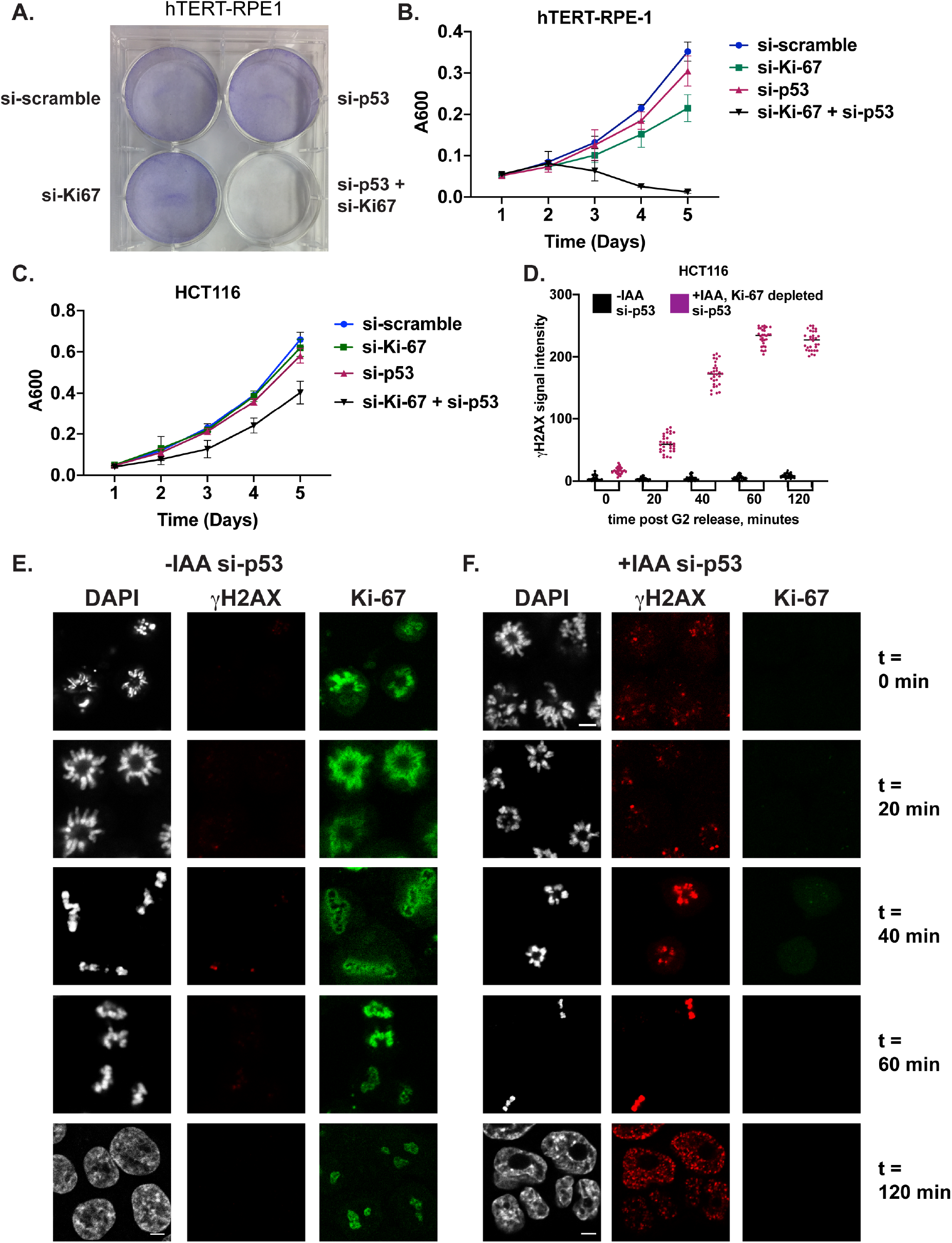
Synergistic effects of co-depletion of Ki-67 and p53. Experiments in this figure were performed using si-p53 (s606), companion experiments using the alternative si-p53 (s607) are shown in Figure S3. (A) Crystal violet staining of hTERT-RPE1 cells treated with the indicated siRNAs. (B) Proliferation of hTERT-RPE1 cells treated with the indicated siRNAs measured with Alamar Blue. Data from 8 replicate populations are shown. (C) Proliferation of HCT116 cells treated with the indicated siRNAs, measured as in panel B. (D) As in Figure 3B, γH2AX was analyzed in RO-3306-synchronized HCT116 cells, except here cells were also treated with si-p53. (E-F) IF images of cells treated as in panel (D). As in Figure 3C-D, the fields of the 120-minute images are of slightly different size than the others. Scale bars are 5 μm.

In addition to greater levels of γH2AX, we observed that co-depletion of Ki-67 and p53 increased the production of structures resembling anaphase bridges (DNA stretched between both segregating chromosome masses) (Figure 6A-B). Anaphase bridges arise from telomeric fusions or from misrepair of DNA damage, and are increased upon exposure to a wide variety of genotoxic agents (39). Observation of anaphase bridges is significant because they contribute to aneuploidy, a very frequent feature of human tumors (40). We observed other anaphase defects with a variety of appearances, in some cases including separated DNA masses resembling lagging chromosomes (Supplemental Figure S5E). We quantified the appearance of anaphase defects in RO-3306-synchronized mitoses in HCT116 cells, either with or without IAA-driven Ki-67 degradation, and either with or without p53 depletion. We observed that anaphase defects were by far most abundant in doubly-depleted cells (Fig. 6A), and that they began to appear after 60 minutes post release, around the time of sister chromosome separation at anaphase (Fig. 5E-F, 6B). These observations suggested that significant genome instability is triggered upon co-depletion of Ki-67 and p53.

**Figure 6.**
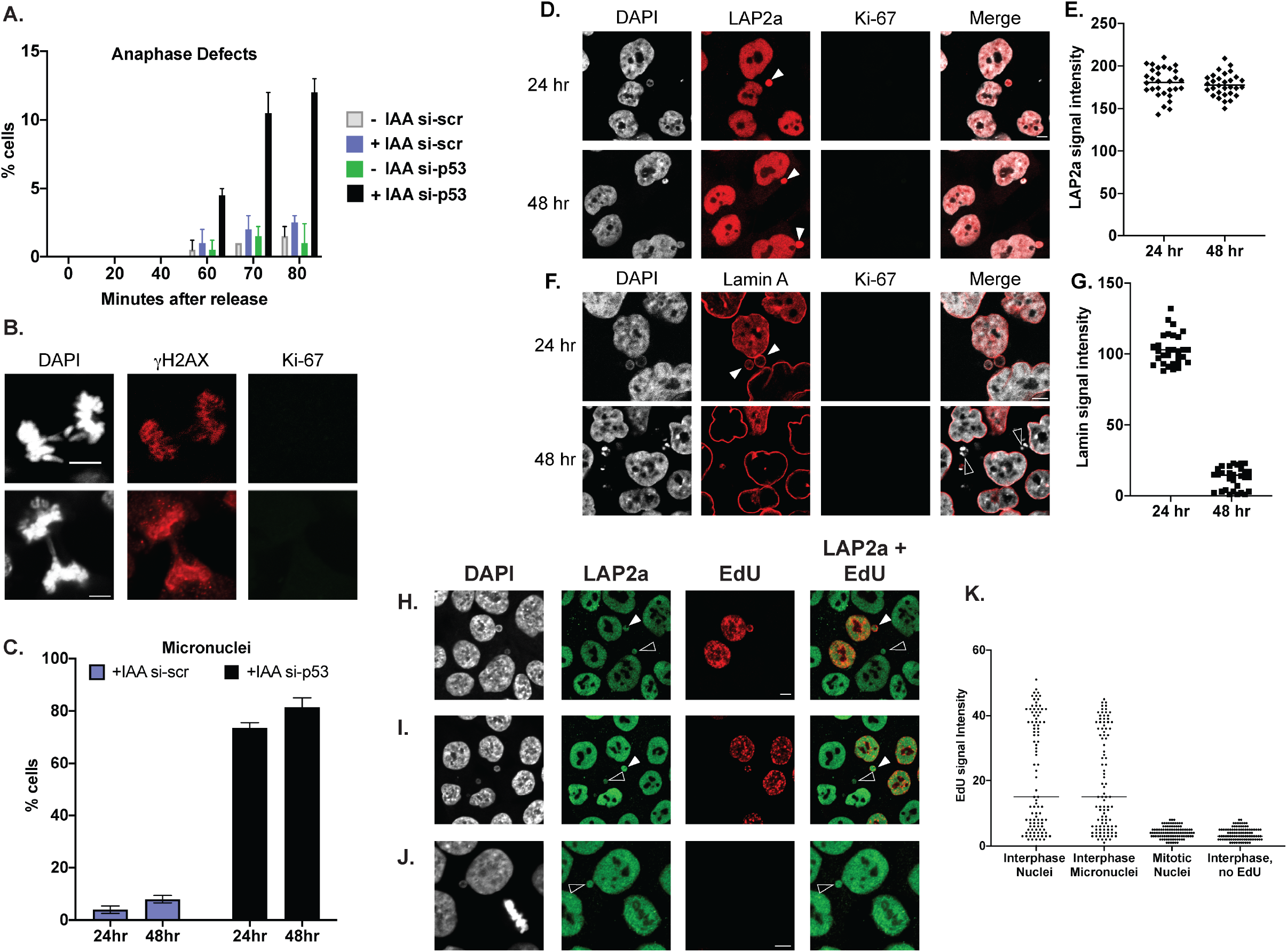
Co-depletion of Ki-67 and p53 results increases the frequency of anaphase defects and micronuclei. (A) Quantitation of defects (bridges and lagging chromosomes) in RO-3306-synchronized HCT116 cells. Note that cells entering anaphase were first observed at the 60-minute time point (Fig. 5E-F). (B) Examples of anaphase defects from RO-3306-synchronized HCT116 cells treated with IAA + si-p53, from the 60-minute point. Additional images are shown in Supplemental Figure S5E. (C) Quantitation of micronuclei in HCT116 cells 24 or 48 hours after the indicated treatments. (D) IF analysis Lap2a in si-p53-treated cells for the indicated times after IAA treatment. (E) Quantitation comparing Lap2a staining from panel D. (F) As in panel D, IF analysis Lamin A in +IAA si-p53 cells. (G) Quantitation of Lamin A staining from panel F. (H-J) HCT116 cells were treated with si-p53 for 72 hours, IAA for the final 24 hours and then labeled with EdU for 20 minutes prior to fixation. K. Quantitation of the EdU intensities of cells illustrated in panels H-J. Interphase cells, micronuclei observed in interphase cells and mitotic cells were quantified as separate classes. Interphase cells that were not pulsed with EdU were quantified as a negative control for background fluorescent signals. Field sizes are the same within panels and for panels H and I. Scale bars are 5 μm.

Indeed, in addition to anaphase defects, microscopic examination indicated additional abnormal nuclear morphologies upon co-depletion of Ki-67 and p53. After the synchronized cells passed through mitosis in the absence of both Ki-67 and p53, these included grossly altered nuclear morphologies and strongly DAPI-stained puncta (Fig. 5F, 120 minute time point). The doubly-depleted cells also displayed small DAPI-stained bodies that appeared to be micronuclei (Fig. 6C-J). Previous studies have shown that anaphase DNA bridges can lead to formation of micronuclei (39, 41), which are small bodies containing single chromosomes or chromosome fragements enclosed within nuclear envelope-derived membranes (42). To confirm that these are micronuclei, we showed that these stained with antibodies recognizing LAP2a (Fig. 6D-E, 6H-J), a protein constitutively found in micronuclei (27). We also observed that after 24 hours of auxin treatment, the micronuclei contained lamin A/C protein around their membrane, but that after 48 hours these levels were greatly reduced (Fig. 6F-G). Such loss of lamin is characteristic of micronuclear envelope degradation (26, 27), a process by which DNA in micronuclei loses nuclear membrane integrity and becomes exposed to inappropriate action of DNA recombination and replication enzymes leading to genome rearrangements. These data reinforce the importance of Ki-67 in genome stability, and suggest that anaphase bridges formed during the first mitosis after Ki-67 loss later become micronuclei. Similar genomic instability cascades are well-documented in the formation of anuploid cells (43).

Recent studies have indicated that defects in DNA replication or broken chromosome bridges formed during interphase can result in aberrant DNA synthesis during subsequent mitoses, on a path towards high levels of genome instability (44, 45). We did not detect DNA synthesis in Ki-67-depleted mitotic cells (Figure 2). However, given our findings regarding p53 (Figure 5), we also tested for DNA synthesis in mitotic cells after acute codepletion of Ki-67 and p53. We labeled cells with EdU to detect DNA synthesis and also stained for LAP2a to enhance detection of micronuclei (Figure 6H-J). We observed frequent EdU labeling of interphase cells in this asynchronous population, with a similar percentage of EdU-positive micronuclei. In contrast, no EdU-positive mitotic cells were detected (Fig. 6I). These data suggest that unscheduled DNA synthesis during mitosis is not a prominent outcome of Ki-67 depletion, either with or without co-depletion of p53. These data also re-enforce our conclusion that mitosis is the critical period for genome protection by Ki-67.

### Distinct effects of acute p53 depletion versus a pre-existing gene deletion

Several recent studies have shown that the presence of p53 can dramatically alter the outcome of synthetic lethality screens (46–48). These and other studies have provided lists of genes that are synthetically lethal with p53, and we note that the *MKI67* gene encoding Ki-67 has not been found among these. We therefore wondered whether loss of Ki-67 in cells that already lack p53 would be cause similarly severe phenotypes. To test this idea, we depleted Ki-67 in a derivative of hTERT-RPE-1 cells in which both *TP53* alleles encoding p53 were deleted via CRISPR (49). As expected, treatment of these cells with si-p53 had no effect. Notably, the p53-null version of RPE-1 cells were not killed by depletion of Ki-67 (Fig. 7A-B). Furthermore, unlike the wild-type RPE-1 cells, these p53^−/−^ cells did not display increased levels of DNA damage upon depletion of Ki-67, as measured via either immunofluorescence or FACS analyses (Fig. 7C). Therefore, the genetic environment can strongly affect the results of Ki-67 depletion. We hypothesize that during the serial passaging of the p53^−/−^ cells in the course of selection and screening, there has been adaptation that overrides the genome instability that would otherwise be caused by loss of Ki-67.

**Figure 7.**
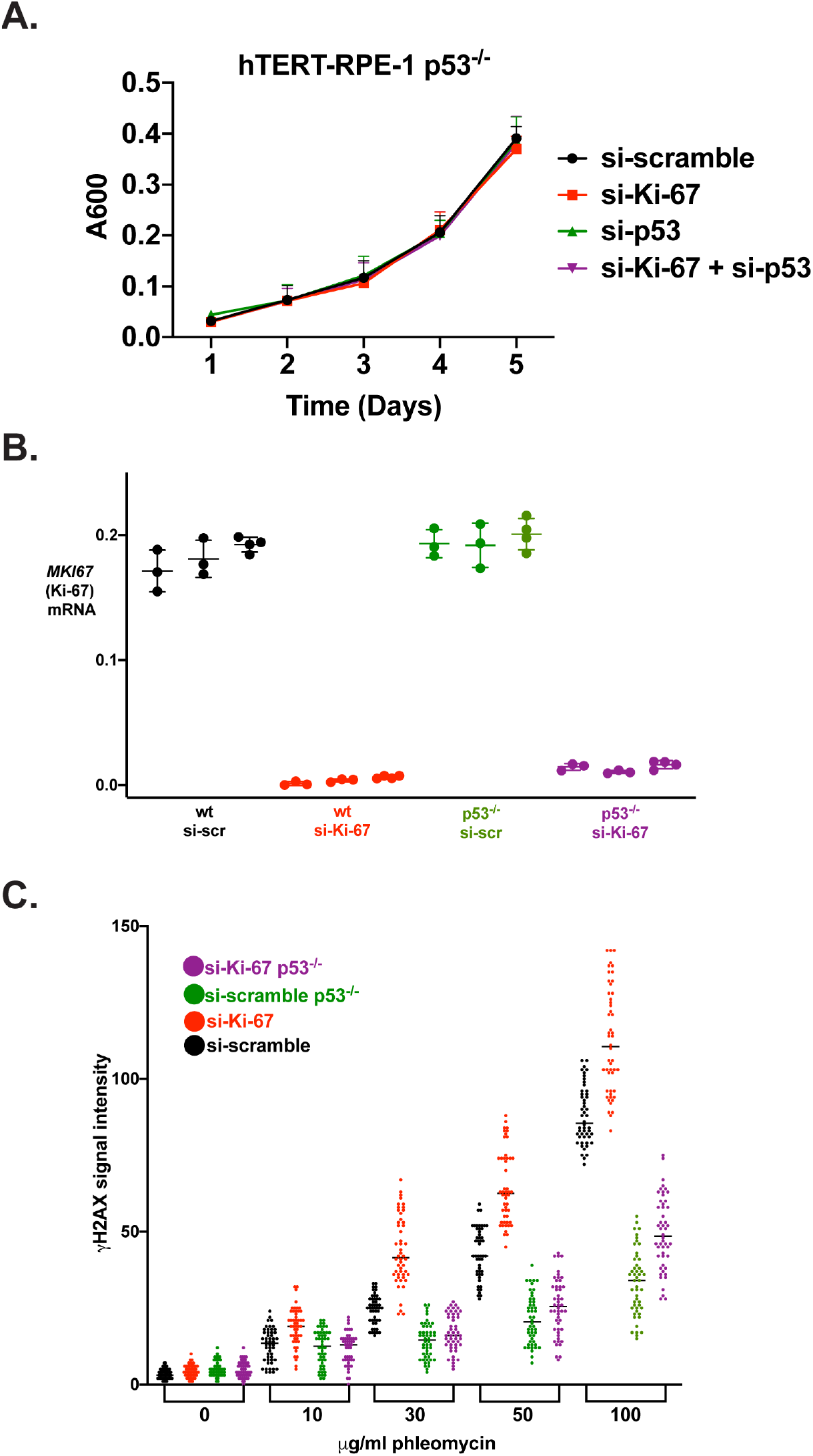
hTERT-RPE1 p53^−/−^ cells do not display growth and DNA damage phenotypes upon Ki-67 depletion. (A) Growth curves of p53^−/−^ hTERT-RPE1 cells treated as indicated, analyzed as in Figure 5B. (B) Validation of Ki-67 depletion by siRNAs in wt and p53^−/−^ hTERT-RPE1 cells. (C) Measurements of γH2AX staining intensity based on IF. Indicated cells were treated with phleomycin to exacerbate damage phenotypes as in Figure 1B.

## Discussion

### Acute depletion of Ki-67 causes DNA damage that originates during mitosis

The role of Ki-67 in genome protection was most evident in mitotic cells (Figure 2), and was evident in the first mitosis after rapid depletion of the protein in the preceding G2 phase (Figure 3). These data suggest that the damage observed in interphase cells originated during mitosis, and that the triggering events are not left over from unrepaired breaks or errors in a previous S phase. Rather, the rapid appearance of damage during progression through mitosis after acute depletion of Ki-67 (Figure 3) leads us to favor the idea that Ki-67 directly protects mitotic chromosomes. In this view, the appearance of γH2AX is one more aspect of defective chromatin maintenance upon Ki-67 depletion, consistent with previously described structural deformations of heterochromatin (11), loss of association and modification of heterochromatin at the nucleolar periphery (10–13), and reduced transcriptional silencing of pericentromeric alpha-satellite repeats (50).

We observed γH2AX and 53BP1 foci in interphase cells upon acute depletion of Ki-67 (Fig. 1). These data suggest that DNA damage explains why checkpoint-proficient hTERT-RPE1 cells acutely depleted of Ki-67 trigger p21-dependent cell cycle delays and transcriptional responses (13). We note that damage was observed in both hTERT-RPE1 cells that induce p21 in response to Ki-67 depletion, and in HCT116 cells that do not display this response (Fig. 1). Therefore, the observed DNA damage is not dependent on the checkpoint-mediated transcriptional signature observed in Ki-67-depleted hTERT-RPE1 cells (13). Consistent with this conclusion, we observed that HCT116 cells display significantly altered levels of very few mRNAs upon Ki-67 depletion (Supplemental Figure S1).

### The C-terminus of Ki-67 protects chromosomes from damage

Ki-67 is a very large protein (over 3000 amino acids), and previous studies have characterized several functional subdomains (Fig. 4C; reviewed in (1)). At the N-terminus is a phosphopeptide-binding FHA domain that interacts with the proteins NIFK and Hklp2 (51, 52). These proteins have been implicated in cancer progression (53) and spindle function (54), respectively, although how these activities may be related to Ki-67 function is not fully clear. Ki-67 also contains a distinct form of a Protein Phosphatase 1 (PP1)-binding site that is also found in the mitotic exit regulatory protein RepoMan (10); this binding site contributes to efficient removal of mitotic phosphorylation from Ki-67 (14). The majority of Ki-67 is composed of an internally repeated set of sixteen ~100 amino domains that contain a site for mitotic CDK1 phosphorylation (14, 55). Finally, at the C-terminus is a leucine-arginine rich “LR” domain (56) that binds DNA in vitro (38) and is required for chromosome binding in cells (15, 20). We show that this C-terminal LR domain is necessary and sufficient for protecting chromosomes from damage during mitosis, either in the presence or absence of p53. This suggests that proteins interacting with Ki-67 domains other than the LR are unlikely to have direct roles in mitotic genome protection. Additionally, these results suggest that the high degree of electrostatic charge across the Ki-67 protein, which is important for maintaining the rod-like shape of individual chromosome arms during mitosis (20, 21) is not required for protection from damage. It remains to be determined whether the Ki-67 C-terminus protects chromosomes from damage during mitosis via steric occlusion of enzymes that can inappropriately break DNA strands, or instead physically stabilizes chromosomes in another manner.

### What is the functional contribution of p53?

p53 is a transcription factor that activates broad responses to DNA damage and other stresses (25, 57). Notably, p53 is mutated in more than half of human cancers (58), and impairing genes that are synthetically lethal with mutated p53 (59) is considered an important path towards therapeutic targets (60). Although most functional contributions of p53 include its activity as a transcription factor, it is clear that the lack of induction of p21 cannot alone account for the Ki-67/p53 synergy, because co-depletion of Ki-67 and p21 in hTERT-RPE1 cells results in loss of G1 checkpoint activation but not synthetic lethality (13). Therefore, a transcriptional role for p53 would have to be p21-independent. However, the rapid kinetics of the appearance of damage leads us to favor a more direct role for p53 in sensing and responding to defects caused by Ki-67 depletion.

Which p53 functions may be most relevant to our observations? We note that chromosome missegregation elevates p53 levels, and that p53 limits proliferation of anuploid cells, contributing to p53’s role as a tumor suppressor (61). Therefore, one possibility is that p53’s role in monitoring chromosome segregation contributes to our observations. For example, depletion of Ki-67 could result in defective chromosome-kinetochore attachements. In the presence of p53, poor attachments may be recognized and fixed, but in its absence, the observed anaphase bridges may result. Alternatively, DNA strand breaks caused by Ki-67 depletion could be catastrophic in the absence of p53. Kinetochore problems and DNA strand breaks could be interrelated, but whether one or the other is the initiating event remains to be explored.

It is clear that the requirement for Ki-67 to protect cells is much reduced in cells with a CRISPR-engineered p53 deletion (Fig. 7) compared to cells experiencing acute depletion of p53 (Fig. 5). These data suggest that cells adapt to the p53 deletion in a manner that compensates for loss of Ki-67. We hypothesize that this is achieved via altered gene expression and are testing this idea now. We also hypothesize that such adaptation mechanisms could explain apparently contradictory genetic studies in the mouse system, where expression of Ki-67 appears to be non-essential for organismal development (11). Another possibility is that not all of the functions of Ki-67 may be shared between humans and mice.

In conclusion, these studies add to the growing list of important mitotic functions for the human Ki-67 protein. Specifically, we show that the C-terminal chromosome-binding domain of Ki-67 protects human cell DNA from damage during mitotic progression. Ki-67 also is the keystone of the mitotic perichromosomal layer (PCL) (10), ensures that mitotic chromosomal arms maintain physical separation (20, 21), and promotes chromosome clustering during mitotic exit, thereby preventing retention of cytoplasmic contents in reforming nuclei (22). It will be of great interest to determine whether these mitotic activities are related to Ki-67’s contributions to the organization and silencing of interphase heterochromatin (10–13).

## Acknowledgements

We thank Judith Sharp and Sharon Cantor for helpful comments on the manuscript. These studies were funded by grant R35 GM127035 from the National Institutes of Health to PDK. We thank Judith Sharp and Michael Blower for the hTERT-RPE1 cells, Daniel Durocher and Sharon Cantor for the hTERT-RPE1 p53^−/−^ cells, and Masatoshi Takagi for the generous gift of plasmids and the HCT116-Ki-67-mAC cells.

## Materials and Methods

### Cell Culture

hTERT-RPE-1 cells were the same isolate previously studied in the laboratory (13), originally obtained from Judith Sharp and Michael Blower (62). These were cultured in DMEM/F12 1:1 ham (Gibco) cat# 11320082 10% FBS (Avantor seradigm, VWR), and 1% penicillin/streptomycin. hTERT-RPE-1 cells with the p53 gene deleted were a kind gift from Daniel Durocher (49) via Sharon Cantor, UMMS. HCT116-Ki-67-mAC-AID cells were a kind gift from Masatoshi Takagi (21) and were cultured in DMEM (Gibco) cat#11995065, with 10% FBS and 100 units/ml penicillin + 100 μg/ml streptomycin. Cells were incubated in 37 °C, 5% CO_2_, and 95% humidity.

Antibodies, siRNAs and PCR primers are listed in Supplemental Table 2.

#### Cell synchronization

Cells were seeded in 6-well plates containing glass coverslips and grown for 48 hours. Cells were first treated according to a double thymidine block protocol (63) involving two sequential rounds of treatment with 2 mM thymidine in complete culture medium for 18hr, followed by washing with phosphate-buffered saline (PBS) and release into drug-free media for 8 hours. Cells were then treated with 10 μM CDK1 inhibitor RO-3306 (Sigma-Aldrich cat. #SML0569, (37, 64)) for 18 hours. For some samples, 0.5 mM indole acetic acid (IAA, Sigma-Aldrich cat. #I5148, (21)) was added for the last 6 hours of RO-3306 treatment to induce degradation of Ki-67 (21). To release cells from the RO-3306 block, they were washed three times with PBS, given fresh drug-free media, and then harvested at the indicated time points.

#### Assay for Ki-67 transgene function

Cells were seeded in 60 mm culture dishes, grown for 48 hours, and then transfected with plasmids as indicated. After 12 hours, some cells were treated with 0.5 mM 3-indole acidic acid (IAA) for 24 hours, and then at 36 hours post-transfection all cells were washed with PBS, harvested by trypsinization, and fixed in 70% ethanol in rotating tubes for 20 minutes. Cells were blocked for 1 hour at 4°C using 1% bovine serum albumin (BSA) in PBS + 0.1% Triton X-100, then primary antibodies (1 ml diluted 1:1000 in blocking buffer) were added for incubation overnight in 1.5 ml microfuge tubes on a rotator at 4°C. Cells were then washed 3 times with PBS + 0.1% Triton X-100, collecting cells by centrifugation for 1 minute at 2000 x g and removing the supernatant by aspiration. Secondary fluorescent antibodies (1:2000 in blocking buffer) were incubated for 1 hour in a tube rotator in room temperature. Cells were washed 3 times with PBS + 0.1% Triton X-100, labelled with 1 μM DAPI diluted in PBS, and washed twice with PBS + 1% BSA + 0.1% Triton X-100 prior to analysis by flow cytometry.

#### Flow cytometry

Labeled cells were filtered through 37μm nylon mesh to enrich for single cells and analyzed in in the UMMS flow cytometry core on a BD LSR II flow cytometer, using wavelengths of 405 nm for DAPI, 488 nm for mClover-tagged Ki-67, and 647 nm for Alexa Fluor 647-labeled secondary antibodies (for γH2AX). 10,000 cells were analyzed for each sample. FCS files were obtained and analyzed using Flowjo 10.6.2.

AMNIS FACS was performed using a Flowsight image cytometer (Luminex) with a 20x AT 0.6 NA objective with 1um pixel size. Images were analyzed using Ideas 6.0 software.

#### Transfection

Plasmids encoding GFP-tagged Ki-67 transgenes were a kind gift from Masatoshi Takagi (15). DNA was purified for transfection using Zymopure midiprep kits (Zymo Research). Plasmid transfections were accomplished using Lipofectamine 2000 (Invitrogen, Inc). 12 hours after plasmid transfections, HCT116 cells were either untreated or treated for 24 hours with 0.5 mM IAA, and then analyzed by flow cytometry or immunofluorescence.

siRNA transfections were performed using Lipofectamine RNAiMAX (Invitrogen) as recommended by the manufacturer. siRNA duplexes were purchased from Ambion (Life Technologies). si-Ki-67 was used at a concentration of 20 μM, si-p21 was used at 40 μM, and siRNAs targeting p53 were used at 80 μM. Concentrations of the silencing control siRNA (“scr”) were matched with the experimental duplex for each experiment. hTERT-RPE1 and HCT116 cells were analyzed 72 hours after siRNA transfections, with phleomycin treatments for the last 5 hours where indicated.

#### Immunofluorescence studies

Cells were seeded in tissue culture-treated 6-well plates for 48 hours, then transfected with siRNAs. In some HCT116 experiments, cells were treated with 500 μM indole acetic acid (IAA) for 24 hours prior to harvest to deplete Ki-67. 72 hours after transfection, cells were washed twice with PBS, and fixed in 4% paraformaldehyde diluted in PBS for 10 minutes at room temperature. Cells were then washed twice with PBS, treated with ice-cold methanol for 20 minutes at −20°C, and blocked for 1 hour at 4°C in PBS + 1% BSA + 0.1% Triton X-100. Antibodies were diluted in the same blocking buffer, added to fixed cells overnight at 4°C. Cells were then washed three times, 5 minutes each with PBS. Secondary fluorescent antibodies were diluted 1:1000 in the same blocking buffer and then added to cells and incubated for 1 hour at room temperature in a dark container. Cells were washed three times with PBS for 5 minutes each, then counterstained with 1 μM DAPI for 1 minute. Cells were washed twice with PBS, and the coverslips were mounted on using Prolong gold antifade (Invitrogen cat. #P36934). Nuclei were marked as regions of interest and average signal intensities were measured using Zeiss Zen Blue software for each analyzed nucleus. IF intensity measurement statistics were analyzed using GraphPad Prism.

#### Visualization of EdU-labeled nascent DNA

HCT116 cells were grown on glass coverslips in DMEM medium as described above. 5-Ethynyl-2-deoxyuridine (EdU, Sigma cat. T511285-5MG) was added to the culture medium at 10 μM for 30 min. After labeling, cells were washed three times with PBS. Cells were fixed in 4% formaldehyde for 20 min. Cells were then rinsed twice with PBS + 0.1 % Triton X-100 and then incubated for 30 min in 100 mM Tris-HCl, pH 8.5, 1 mM CuSO_4_, 100 mM ascorbic acid, and 50 mM MB-Fluor 595 azide (Click Chemistry Tools, cat. 1169-5) for click-chemistry labeling (13, 65). After staining, the cells on coverslips were washed three times with PBS + 0.5% Triton X-100 for 5 min each. Cells were then counterstained with DAPI, mounted onto microscope slides, and imaged by fluorescence microscopy as described above.

#### RNA isolation and real-time quantitative PCR

Total RNA was isolated from cells at 72h post-transfection using Qiazol (Qiagen) following the manufacturer’s instructions and purified using an RNeasy kit (Qiagen). One microgram of RNA was subjected to reverse transcription with SuperScript II reverse transcriptase (Invitrogen). qPCR reactions were performed on an Applied Biosystems StepOnePlus machine (Life Technologies), using Fast Sybr mix (Kapa Biosystems). The program used was as follows: hold at 98°C for 30 s followed by 40 cycles of 95°C for 10 s and 60°C for 30 s. All the signals were normalized to that of beta-actin (loading control) and the 2^−ΔΔ^*CT* analysis method was used for quantification (Life Technologies). Primer sequences were designed by use of Primer3Plus software. All oligonucleotides for qPCR are listed in Supplemental Table 2.

#### RNA-seq

For each biological replicate sample, 1 x 10^6^ HCT116 cells were seeded into each of three 35 mm dishes, grown for 2 days and then untreated or treated with 0.5 mM IAA for 24 hours. Media was removed, 0.7 ml Qiazol (Qiagen) was added to each dish and the lysate was collected into Qiashredder tubes (Qiagen) and centrifuged at 12,000 x g for 30 seconds for homogenization. Total RNA was extracted using RNeasy kits (Qiagen) as recommended by the manufacturer. Total RNA concentrations were measured using Qubit reagents (Invitrogen). mRNA was enriched on oligo-dT beads, and reverse transcribed and sequenced using a PE150 protocol with Illumina reagents at the Novogene corporation.

Paired-end reads were aligned to human primary genome hg38, with star_2.5.3a (66), annotated with GENCODE GRCh38.p12 annotation release 29 (67). Aligned exon fragments with mapping quality higher than 20 were counted toward gene expression with featureCounts_1.5.2 (68). Differential expression (DE) analysis was performed with DESeq2_1.20.0 (69). Within DE analysis, ‘ashr’ was used to create log2 Fold Change (LFC) shrinkage for each comparison (70). Significant DE genes (DEGs) were identified with the criteria FDR < 0.05.

#### Parameters

**Genome:** hg38.primary.fa

**GTF:** gencode.v29.primary_assembly.annotation.gtf

#### Star parameters

STAR --runThreadN {threads} \
--genomeDir {input.index} \
--sjdbGTFfile {input.gtf} \
--readFilesCommand zcat \
--readFilesIn {params.reads} \
--outFileNamePrefix mapped_reads/{wildcards.sample}. \
--outFilterType BySJout \
--outMultimapperOrder Random \
--outFilterMultimapNmax 200 \
--alignSJoverhangMin 8 \
--alignSJDBoverhangMin 3 \
--outFilterMismatchNmax 999 \
--outFilterMismatchNoverReadLmax 0.05 \
--alignIntronMin 20 \
--alignIntronMax 1000000 \
--alignMatesGapMax 1000000 \
--outFilterIntronMotifs RemoveNoncanonicalUnannotated \
--outSAMstrandField None \
--outSAMtype BAM Unsorted \

#### Imaging

Images were acquired using a Zeiss LSM700 confocal microscope equipped with laser lines of 405/488/594/647nm and suitable emission filters. A Plan-Apochromat 63x/1.40 Oil DIC m27 objective was used. For analysis of 53BP1 foci, a Zeiss Axio-observer Epifluorescence microscope with mounted Axiocam 506 monochrome camera and automated stage was used. Data were analyzed using Zeiss Zen blue software.

#### Proliferation Assay

Five thousand cells were seeded into 96-well plates in 200 μl of complete culture medium and transfected with siRNAs as described above. For each time point starting at 24 hours after transfection, fresh culture media with a 1:100 dilution of Alamar blue (Bio-Rad cat. #BUF012A) was then added to the cells for 3 hours. Viable cells reduce resazurin (Alamar blue) to resorufin which was measured by absorbance at 600 nm. Absorbances were measured using a Glomax (Promega) plate reader every 24 hours for 5 days.

## Supplemental Figures

**Supplemental Figure S1.**
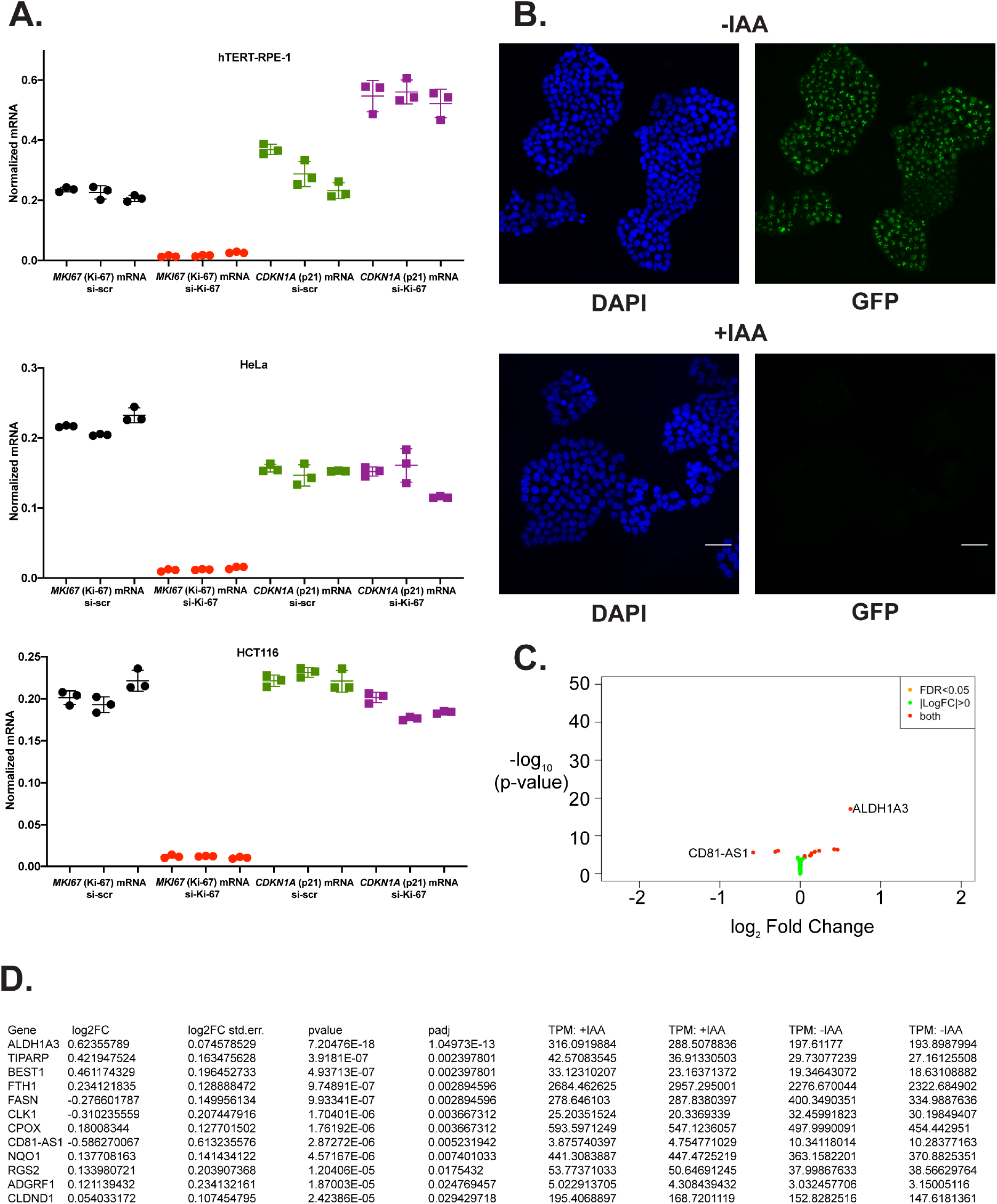
Effects of Ki-67 depletion on steady-state mRNA levels. A. qRT-PCR analysis of *MKI67* (round symbols) and *CDKN1A* (square symbols) mRNAs, encoding Ki-67 and p21, respectively. The indicated cell lines were transfected with the indicated siRNAs. As previously reported, depletion of Ki-67 induced p21 mRNA in RPE-1 but not HeLa cells (13), and we show here that HCT116 cells also lack this checkpoint response. Ki-67 mRNA levels confirm the expected effect of the Ki-67-targeted siRNAs. In each case, actin-normalized mRNA levels are reported for triplicate technical measurements from 3 independent biological replicate experiments. B. Images of cells used for the RNA-seq experiment, either untreated (-IAA) or auxin-treated (+IAA). The GFP channel detects the mClover-tagged Ki-67 protein. Scale bar 50 μm. C. Volcano plot of RNA-seq data for HCT116-Ki-67-mAC cells treated or untreated with auxin (IAA) to deplete Ki-67. The twelve genes indicated by the red spots are listed in panel D. Note that the greatest log2 fold change value for any gene is 0.62. D. List of the twelve genes significantly altered (adjusted p-value < 0.05, out of 34826 genes measured) upon IAA-mediated degradation of Ki-67 in HCT166 cells. Left to right, the log2 Fold Change, Standard error of the log2 fold change, p-value, adjusted p-value, and raw Transcripts Per Million (TPM) values for the biological replicate experiments are presented. See Supplemental Table 1 for detailed analysis of the RNA-seq data.

**Supplemental Figure S2.**
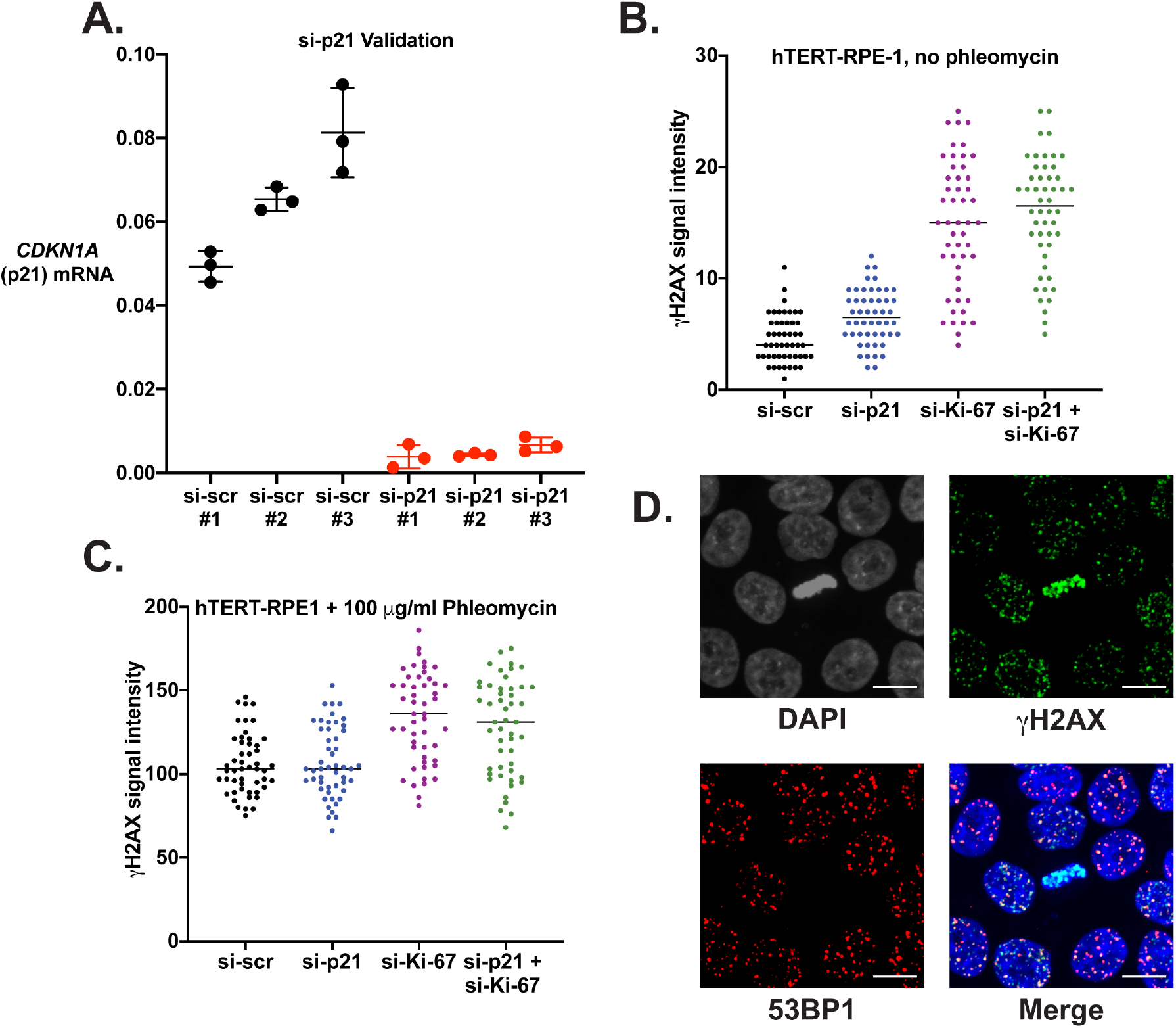
Characterization of p21 depletion phenotypes and 53BP1 foci. (A) RT-PCR validation of si-p21-mediated depletion of p21 mRNA levels in hTERT-RPE1 cells. Triplicate measures of three samples for each condition were analyzed. (B) Depletion of p21 does not exacerbate damage caused by Ki-67 depletion. hTERT-RPE1 cells were treated with the indicated siRNAs and analyzed as in Figure 1B in the absence of phleomycin. (C) As in panel B, except cells were treated in the presence of 100 μg/ml phleomycin. Depletion of p21 did not significantly affect γH2AX levels (Welch’s t test, p = 0.30 and 0.27 in panels B and C, respectively). (D) IAA-treated HCT116 cells were prepared for IF with antibodies recognizing γH2AX and 53BP1. Note the cell in metaphase at the center of the image, which displays γH2AX but not 53BP1 staining.

**Supplemental Figure S3.**
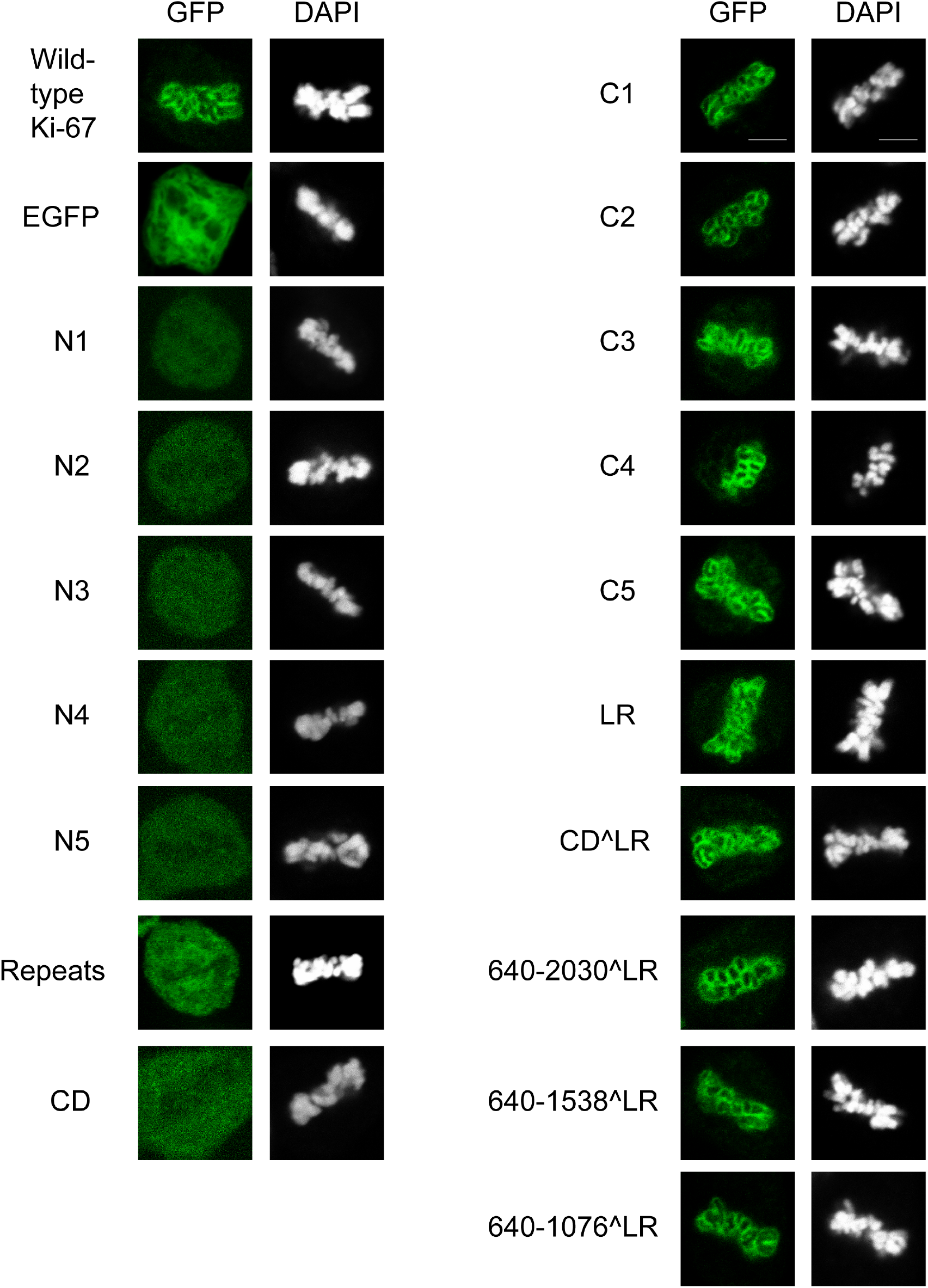
Mitotic localization of transgenes. HCT116 cells were transfected with constructs encoding the indicated Ki-67 transgenes (see Figure 4C for diagram). 12 hours after transfection, endogenous Ki-67 was degraded by addition of 0.5 mM IAA, and cells were analyzed 24 hours later. The GFP-tagged transgene was visualized along with DAPI-stained DNA. Cells expressing full-length, wild-type Ki-67-GFP and the empty vector encoding eGFP are shown on the upper left. The rest of the left-hand column are constructs that did not protect cells from DNA damage. On the right are cells expressing transgenes that did protect from damage. All of the latter class contain the chromosome-binding C-terminal domain, and this imaging confirms the expected mitotic localization on chromosomes.

**Supplemental Figure S4.**
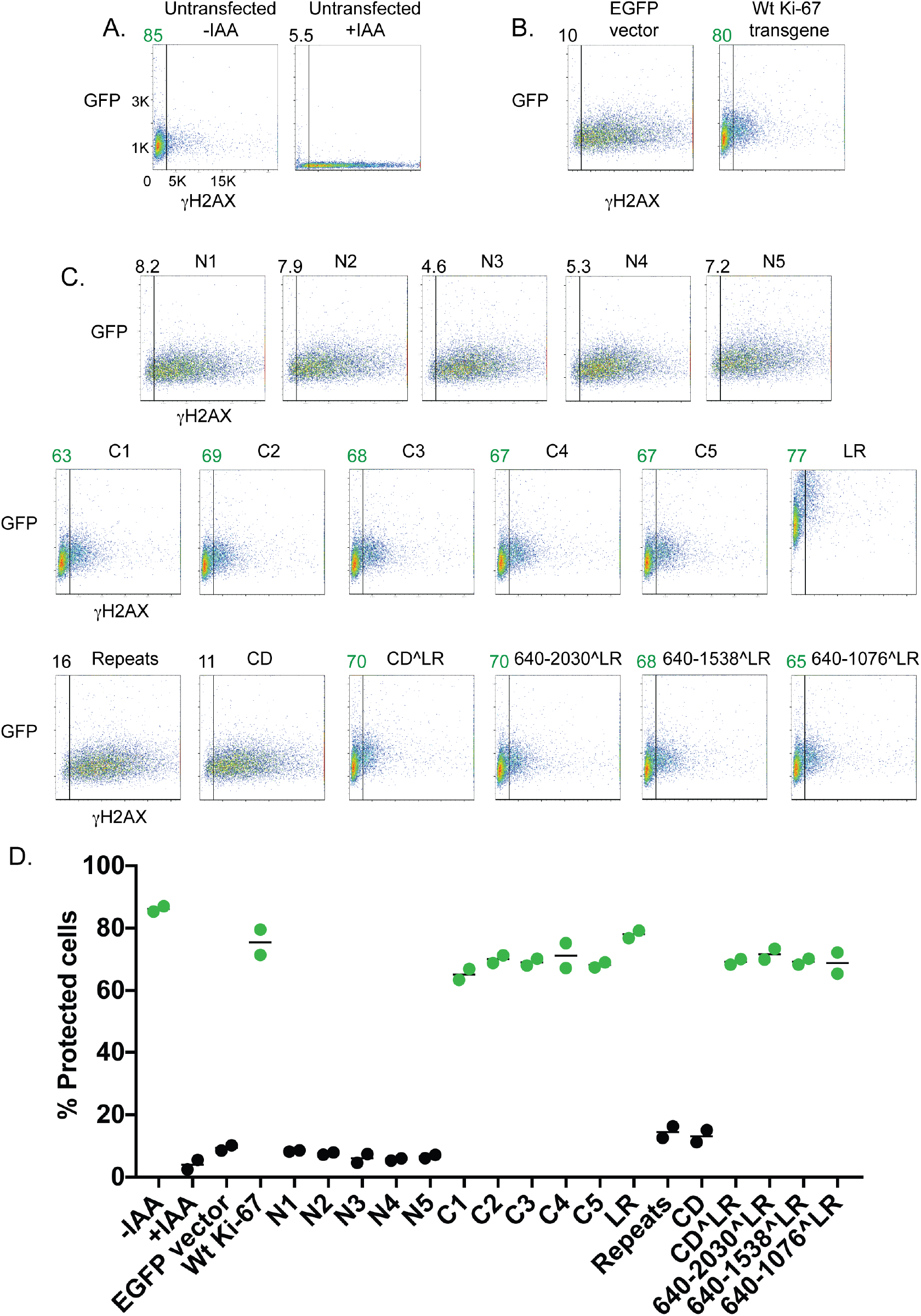
Domain analysis of Ki-67 in cells depleted of p53. Cells were analyzed as in Figure 4, except here cells were also treated with si-p53 (s606). Panel A shows the effect of auxin-mediated Ki-67 depletion in the absence of plasmid transfection. Panel B shows complementation by wild-type Ki-67. Panel C and D show FACS analysis and quantitation as in Figure 4. Note that the magnitude of γH2AX signals (x-axis) in these samples is greater than observed in Figure 4, indicative of the increased damage caused upon co-depletion of p53 and Ki-67. Nevertheless, the C-terminal domain of Ki-67 still protects from this damage.

**Supplemental Figure S5.**
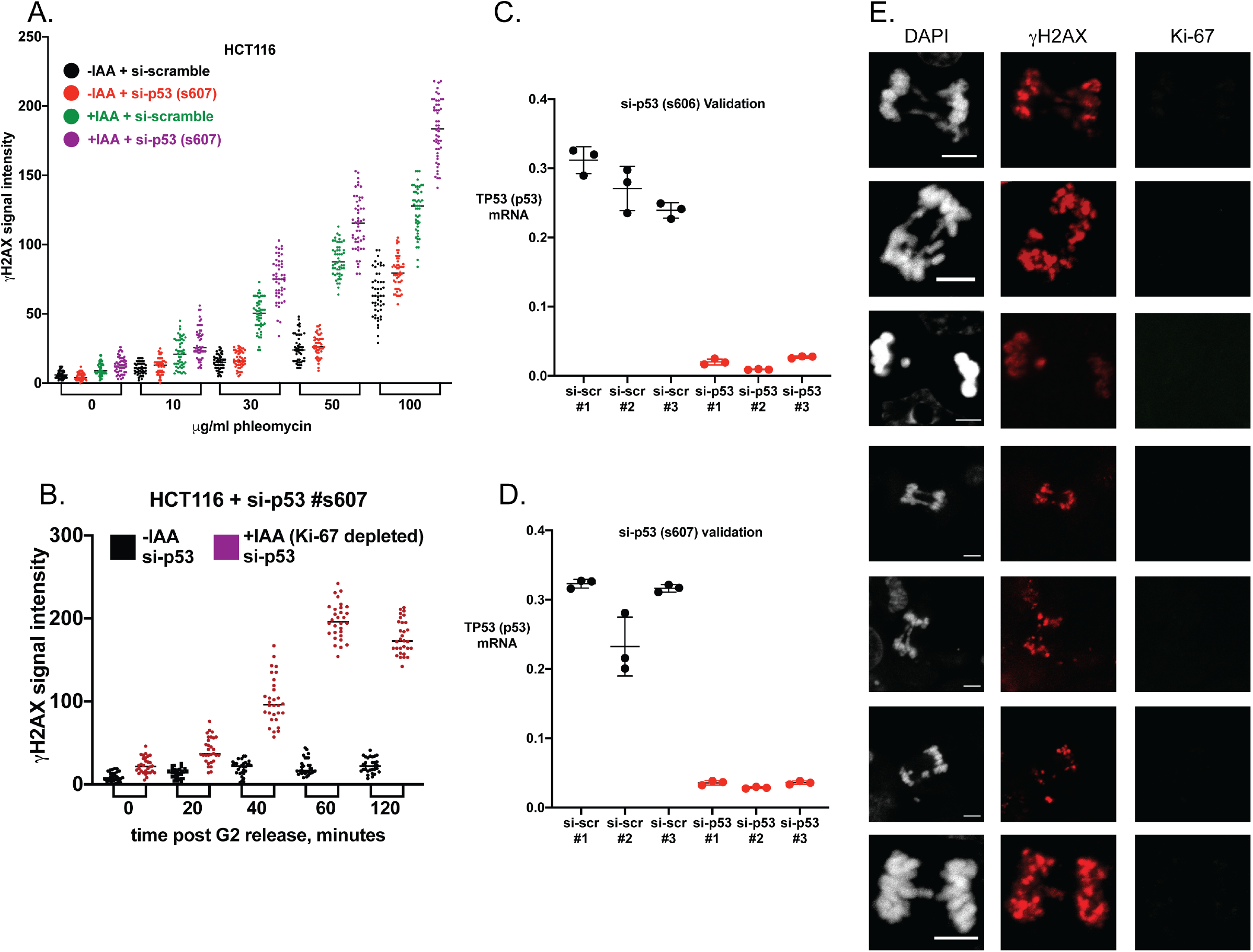
Confirmation of synergistic phenotypes with a second si-p53 reagent (s607). (A) HCT116 cells were treated as indicated and γH2AX levels were analyzed as in Fig. 1D. (B) γH2AX levels in RO3306-synchronized HCT116 were analyzed as in Figure 5D, except that si-p53 s607 was used here rather than s606. (C) Validation of si-p53 (s606) depletion of p53 mRNA levels in HCT116 cells. (D) Validation of si-p53 (s607) depletion of p53 mRNA levels in HCT116 cells. Triplicate measures of three samples for each condition were analyzed. (E) Images of additional anaphase defects as in Figure 6B. All images are from HCT116 cells treated with IAA + si-p53. Cells were all from 60 minutes after release from RO-3306, except the bottom image, which is from 70 minutes after release. Scale bars are 5 μm.

Supplemental Table 1. RNA-seq analysis of HCT116 cells with and without auxin-mediated depletion of Ki-67.

Supplemental Table 2. Antibodies, siRNAs and PCR primers used.

